# A cognitive state transformation model for task-general and task-specific subsystems of the brain connectome

**DOI:** 10.1101/2020.12.23.424176

**Authors:** Kwangsun Yoo, Monica D Rosenberg, Young Hye Kwon, Dustin Scheinost, R Todd Constable, Marvin M Chun

**Author notes:** Correspondence: Dr. Kwangsun Yoo.

## Abstract

The human brain flexibly controls different cognitive behaviors, such as memory and attention, to satisfy contextual demands. Much progress has been made to reveal task-induced modulations in the whole-brain functional connectome, but we still lack a way to model changes in the brain’s functional organization. Here, we present a novel connectome-to-connectome (C2C) state transformation framework that enables us to model the brain’s functional reorganization in response to specific task goals. Using functional magnetic resonance imaging data from the Human Connectome Project, we demonstrate that the C2C model accurately generates an individual’s task-specific connectomes from their task-free connectome with a high degree of specificity across seven different cognitive states. Moreover, the C2C model amplifies behaviorally relevant individual differences in the task-free connectome, thereby improving behavioral predictions. Finally, the C2C model reveals how the connectome reorganizes between cognitive states. Previous studies have reported that task-induced modulation of the brain connectome is domain-specific as well as domain-general, but did not specify how brain systems reconfigure to specific cognitive states. Our observations support the existence of reliable state-specific systems in the brain and indicate that we can quantitatively describe patterns of brain reorganization, common across individuals, in a computational model.

## Introduction

The human brain is versatile, regulating various behavioral and cognitive functions appropriately for different task situations. A fundamental question in neuroscience has been to understand how the brain can flexibly generate such diverse functions. Studies have developed approaches at a wide spectrum of scales, from the micro-scale (e.g., cellular or molecular [Furey et al., 2000; Lisman et al., 2018]) to the macro-scale (whole-brain [Gray et al., 2003; Rosenberg et al., 2016]), to reveal detailed working mechanisms of human cognition, including memory, attention, and decision-making.

The brain supports cognition through the coordinated activity of distributed areas, which is often studied as functional connectivity and the connectome—the whole-brain connectivity network. To accomplish any given cognitive function, it is well understood that multiple brain areas work together, rather than one area operating in isolation. Furthermore, the functional connectome is unique to each individual like their fingerprint (Finn et al., 2015). The connectome retains its individuality irrespective of the type of cognitive involvement, and it can be measured even when not engaged in any explicit task, known as the task-free, resting-state, or intrinsic functional connectome (Buckner et al., 2013; Park and Friston, 2013; Smith et al., 2015).

We do not, however, understand how the brain functionally reorganizes from a task-free state to a task-specific state and from one task state to another. Comparing task-related and task-free connectomes reveals both task-general components such as integrative brain hubs involved in diverse tasks (Cole et al., 2013; Gratton et al., 2016) as well as task-specific differences (Gonzalez-Castillo et al., 2015; Shine et al., 2016). Moreover, task-induced differences can be dominated by group and individual factors, as well as their interactions (Gratton et al., 2018), rendering task effects to be negligible. Despite the recent progress suggesting task-induced modulations on brain activity, we still lack a way to quantitatively describe and model how the functional connectome reorganizes between cognitive states.

To expand our understanding of how the brain supports human cognitive functions at a system level, it would be ideal to develop a brain reconfiguration model that can mathematically transform one state-specific connectome to another. Such a model could, for instance, generate multiple individual task connectomes from a single task-free connectome. Successful modeling of brain network reconfigurations would suggest that cognitive tasks modulate the brain connectome in a reliable and systematic way, and would make the task-free connectome even more informative and useful than previously possible.

Here, we introduce a novel connectome-to-connectome (C2C) state transformation modeling framework that enables us to generate task-specific connectomes from task-free scans (**Figure 1**). We demonstrated our framework using a functional magnetic resonance imaging (fMRI) dataset from the Human Connectome Project (HCP, S1200 data release). We constructed the C2C state transformation model for each of seven brain states defined by seven different tasks (Emotion, Gambling, Language, Social, Relational, Motor, and Working Memory) in the HCP. We demonstrated that the C2C model accurately generates task connectomes at an individual level with high degree of specificity across the seven cognitive states. In addition, the C2C model amplifies behaviorally relevant information of the task-free connectome, thereby improving predictions of individual behavior.

**Figure 1.**
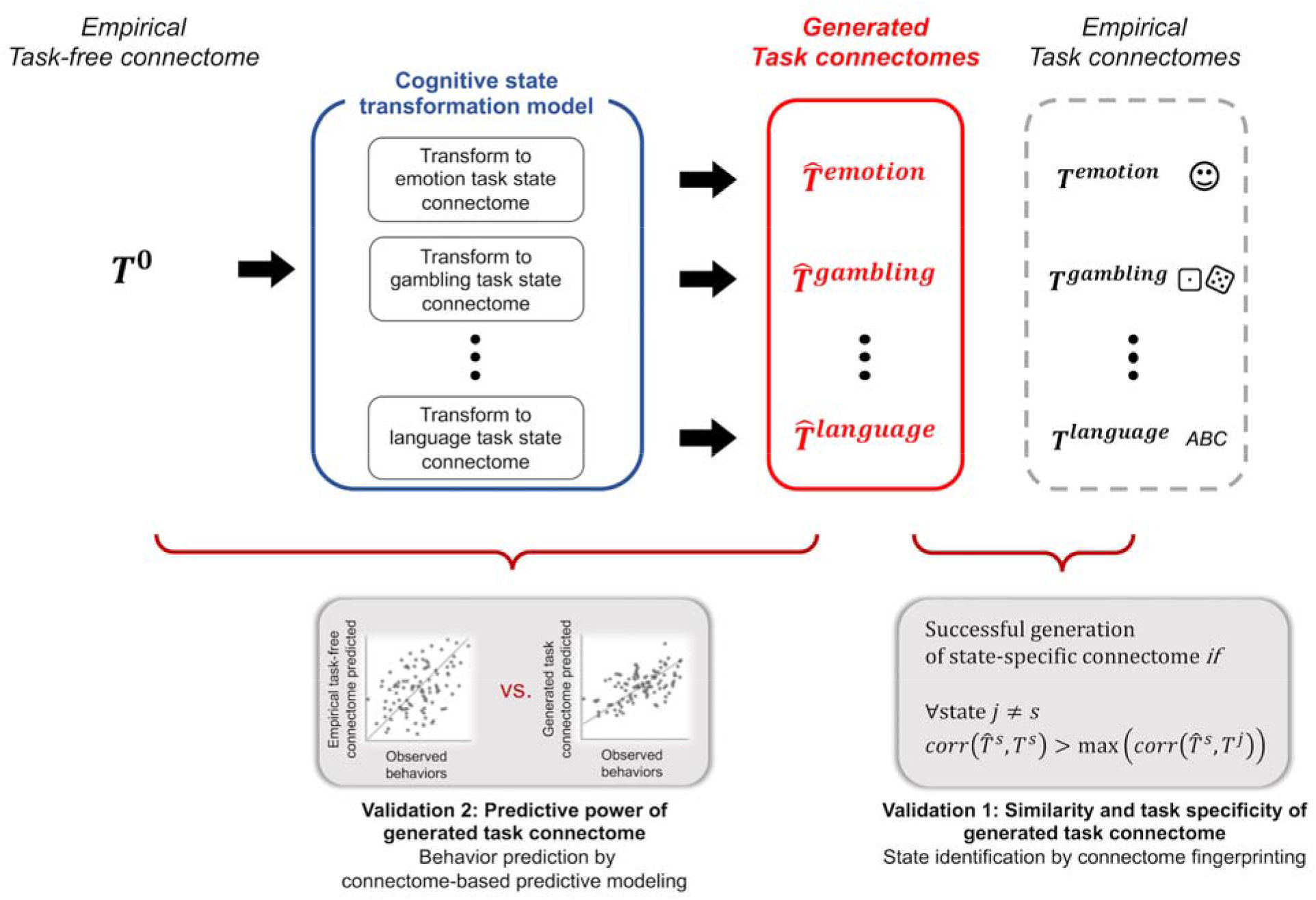
The brain state transformation model and its validation. The connectome-to-connectome (C2C) state transformation model transforms task-free connectome to generate task connectomes. The C2C-generated task connectomes were validated using multiple approaches, demonstrating whether the generated task connectome specifically resembles the empirical connectome of the same task (Validation 1) and whether the generated task connectome better predicts individual behaviors than the observed task-free connectome (Validation 2).

Importantly, the C2C modeling framework provides a way to expand our understanding on the functional mechanism of the large-scale brain network in supporting diverse human cognition. The accurate generation of individual task-specific connectomes by a quantitative model can formulize the brain reorganization pattern in an explainable way. The presented model consists of relatively simple linear functions, *principal component analysis* and *partial least square regression.* This simplicity makes the model transparent and interpretable, quantifying the brain’s functional reorganization in response to specific cognitive goals. We explored the connectome reorganization between cognitive states by scrutinizing the model-defined state transformation. Overall, our observation suggests that a cognitive context introduces a group-common rule of reorganization in functional connectome. This standardized reorganization acts on individuals’ unique functional connectivity patterns, and thereby induces individualized task-specific connectomes.

## Results

### Connectome-to-connectome (C2C) state transformation model accurately generates task-specific connectomes from task-free connectomes

We validated the proposed framework first by testing whether constructed C2C models accurately generate task-specific connectomes. To study this, we assessed the similarity of the model-generated task connectomes with corresponding empirical task connectomes at the whole-brain connectome level as well as at the edge level. At the connectome level, the spatial pattern of the generated task connectomes was significantly correlated with that of the empirical task connectomes (**Figure 2A**). The similarities of the generated connectomes with the corresponding empirical connectomes ranged from *r=0.723* for the WM task state to *r=0.643* for the Emotion task state.

**Figure 2.**
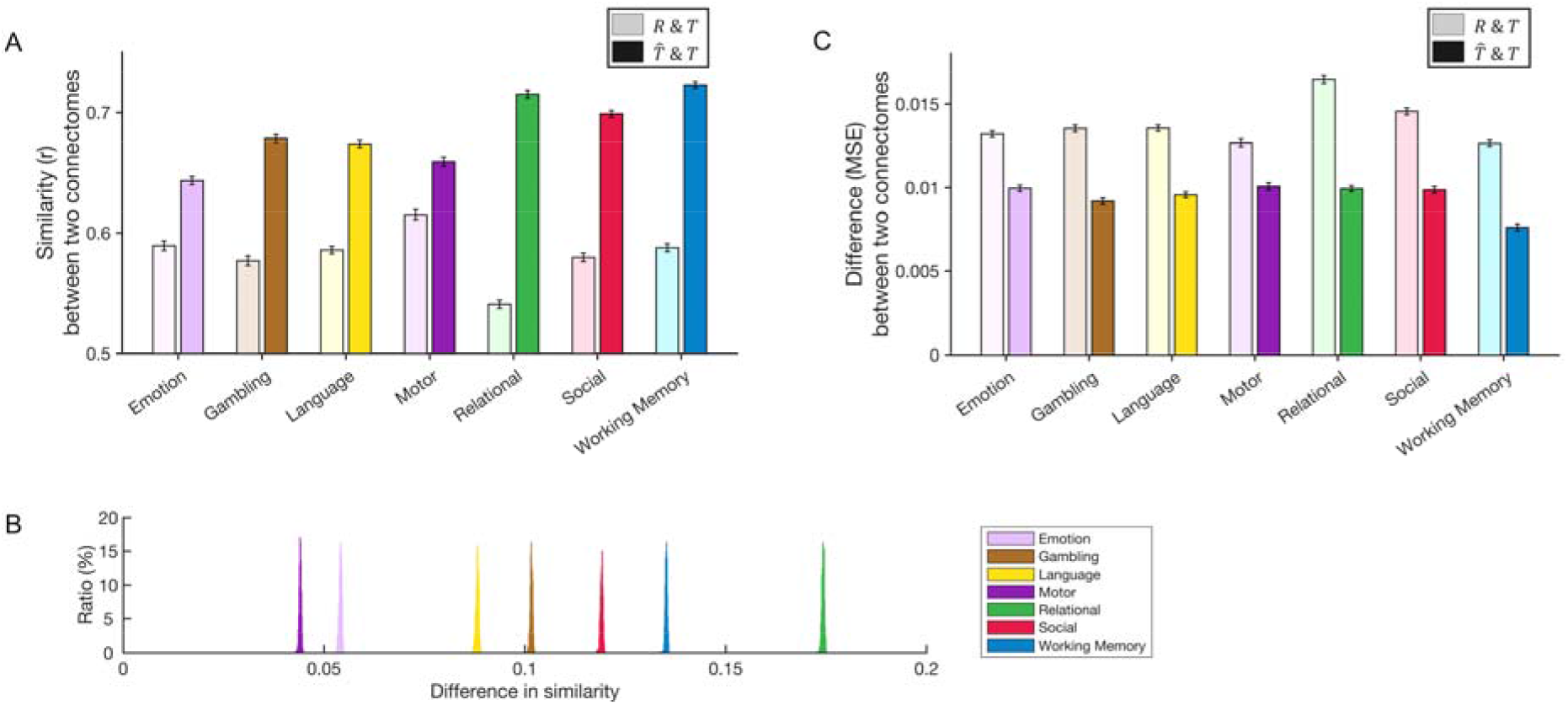
A) The empirical task connectome is more similar to the estimated task connectome than to the task-free connectome in all seven task states. In each task column, a darker bar (right) represents similarity between the estimated and empirical task connectome, and a lighter bar (left) represents a similarity between the empirical task-free and task connectome. Results are from 1,000 iterations of 10-fold cross validation. Error bars represent standard error across 316 subjects. (*R:* observed task-free (rest) connectome, *T:* empirical task connectome, 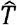. generated task connectome). B) The distribution of difference between these two similarities *(similarity between estimated and empirical task connectomes – similarity between empirical task-free and task connectomes)* across 1,000 iterations. C) Mean square error between estimated and empirical task connectomes is lower than that between observed task and task-free connectomes. Error bar represents standard error across 316 subjects. *(R:* observed task-free (rest) connectome, *T:* empirical task connectome, 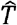 generated task connectome).

Importantly, the similarity between the model-generated and empirical task connectomes is significantly higher than the similarity between the task-free and the empirical task connectomes. We obtained the distribution of differences between these two similarity scores across 1,000 iterations of 10-fold validation. The distribution was fully above the null difference value of 0 (**Figure 2B**), suggesting that the C2C models generate more accurate task-specific connectomes. In addition, the generated Gambling, Language, Relational, Social, and Working Memory connectomes were more similar to their corresponding empirical task connectomes than to the observed task-free connectome (**Figure S1**). This higher similarity of model-generated connectomes suggests that the C2C models meaningfully transformed the task-free connectome to task-specific states.

In order to evaluate the generation accuracy at the edge level, we computed MSE between the model-generated and the empirical task connectomes (**Figure 2C**). We compared MSE of the model-generated connectomes to MSE between empirical task-free and task connectomes. The model-generated connectomes had a reduced MSE compared to the MSE of the empirical task-free connectomes. These results held for all seven task conditions, confirming that the C2C models accurately generated task-specific connectome matrices. In sum, we here demonstrated the successful modeling of the connectome transformation between cognitive states.

We confirmed that noise removal in individual connectomes, by PCA alone, does not explain the increased connectome similarity by the C2C models. The noise removal in the rest connectome rather decreases its similarity to the task connectome for every task state (**Figure S2**). This result indicates that the state transformation in the C2C models, using PLS regression, is essential to accurately estimate task-specific connectomes.

We further tested how the amount of training data influences C2C modeling. We repeated the same 1,000 iterations of 10-fold cross-validation using 50, 100, and 200 subjects. We observed that increasing the number of subjects enabled better C2C models, improving model accuracy in generating task connectomes. Notably, even with only 45 training samples (**Figure S3**), the C2C models generated task connectomes that were significantly more similar than the rest connectome to the empirical connectomes.

### C2C model generates functional connectomes specific to cognitive states

It’s possible that the C2C models are only picking up on task-general differences between task-specific and the task-free states, given domain-general differences in multiple task-specific states relative to the task-free state (Cole et al., 2014). It is important to demonstrate that the C2C models capture task-specific transformations, not just a general transformation from task-free to task-related states. Accordingly, we examined the specificity of the C2C modeling across cognitive tests. We compared all seven model-generated task connectomes with all seven empirical task connectomes. In **Figure 3A**, on-diagonal elements present within-task similarities of the estimated and empirical connectome, and off-diagonal elements show cross-task similarities between them. This similarity matrix was estimated for each subject and then averaged to provide a group-level result. For every task state, within-task similarity was higher than cross-task similarities (**Figure 3A**). For example, the model-generated connectome for the WM task is more similar to the empirical WM task connectome *(r=0.723)* compared to the other empirical task connectomes.

**Figure 3.**
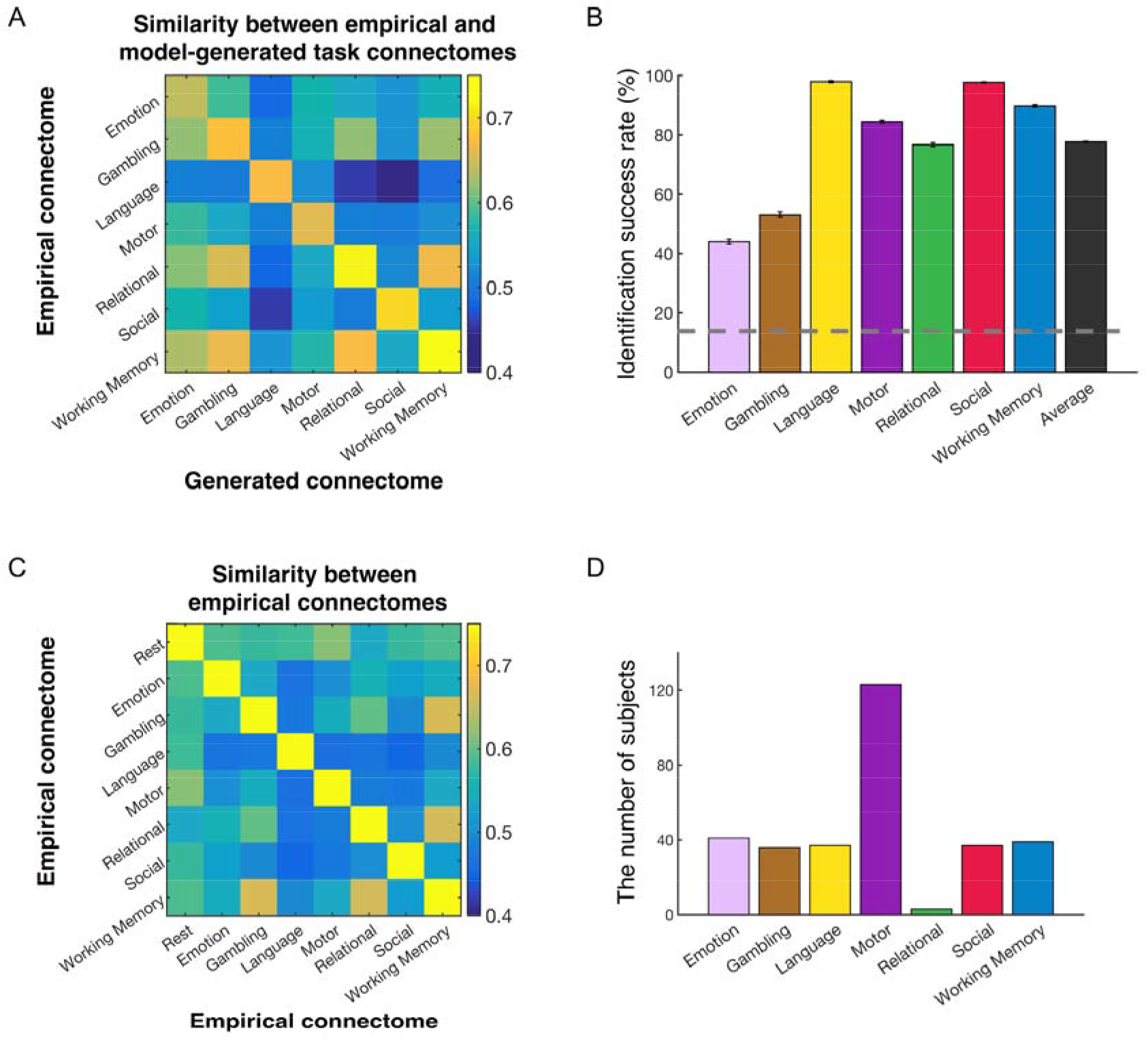
A and B) Task specificity of the model-generated task connectomes. A) Spatial similarity between empirical and generated task connectomes. On-diagonal elements show within-task similarities of the generated and empirical connectomes, and off-diagonal elements represent cross-task similarities. For all task states, within-task similarity was higher than all cross-task similarities. B) Task identification success rate was assessed with a state fingerprinting approach. The generation of a task connectome is considered successful if and only if the generated task connectome maximally resembles the empirical connectome from the same task state compared to those from the other task states. State fingerprinting is more conservative than the spatial similarity analysis shown in A. The gray dashed line represents the chance level of 14.3% (=100/7). Error bar represents standard deviation from 1,000 iterations. C) Spatial similarity of empirical connectomes between one task-free and seven task-specific states. On-diagonal elements show within-task similarities of the empirical connectomes (*r*=1), and off-diagonal elements represent cross-task similarities. D) Classification of empirical task-free connectomes into one of seven task states. The classification was performed in each subject and then individual results were accumulated for the group-level results shown in the bar graph. The *y* axis represents the number of subjects whose task-free connectome was classified as each task-specific connectome.

To better visualize task specificity and the accuracy of C2C-generated task connectivity, we plotted the predicted contrast between task and rest connectomes along with the empirical contrast between task and rest connectomes (**Figure S4**). Figure S4 shows accurate predictions of connectome differences across tasks for four representative individuals, illustrating the explanatory precision of the C2C-generated connectomes, and showing that the model generates individual-specific connectomes rather than population-mean connectomes.

We further assessed task specificity using a connectome fingerprinting approach (Finn et al., 2015), “state fingerprinting”. In this approach, the model-generated task connectome was compared with all seven empirical task connectomes, and the task state of the empirical connectome that exhibited the highest similarity was identified. For example, if a connectome generated by a C2C model constructed for the WM task matched best with the empirical WM task connectome, then this model is considered task-specific. This identification procedure was applied to every single subject for each task. Then for each task state we computed an identification success rate by accumulating individual success or failure (**Figure 3B and S5**). A high identification success rate indicates high task specificity of the C2C modeling framework, and a low rate indicates low task specificity (or task generality). The success rate averaged across the seven tasks was 74%, significantly higher than the chance level of 14.3% (**Figure 3B**). The success rates of all seven task states were significantly higher than chance level as well. Together with the higher within-task similarity demonstrated earlier, the successful identification of task states using the model-generated task connectomes, indicate that the C2C state transformation models provide reliably high task specificity.

We then investigated if the high identification success rate of specific tasks (here, for instance, language [yellow] and social [red] in **Figure 3B**) can be attributed to the similarity of the empirical task-free and the task-specific connectomes (**Figure 3C**). To do this, we analyzed to which task-specific state the task-free connectome has the most similar pattern. This analysis was again performed for every subject and then aggregated to produce a group-level result. We found that in most cases (more than one third of total subjects) the motor task connectome is the most similar to the task-free connectome (**Figure 3D**). All other states but Relational has similar numbers of subjects in which the task-free connectome is the most similar to the target task. This result indicates that the successful identification of cognitive states by the C2C models cannot be explained by the observed similarity between task-free and specific task states, and the C2C models accurately generate the connectome of cognitive states by extracting appropriate transformation between cognitive states.

### C2C model amplifies behaviorally relevant information of individuals

Next, we investigated whether C2C models amplify information unique to individuals, increasingly important for both clinical and research applications. For this, we tested our models in predicting individual intelligence with connectome-based predictive modeling (CPM, Shen et al., 2017). Previous studies reported better behavioral prediction by task-induced connectomes compared to task-free connectomes (e.g., Green et al., 2018; Yoo et al., 2018). In this point, we asked if generating task-specific connectomes improves accuracy of fluid intelligence predictions. The C2C modeling methods would not improve behavioral predictions if this modeling simply adds edge-wise group-averaged differences between the empirical task-free and task-specific states during state transformation. Since CPM is based on a linear regression, adding the same value to all input across subjects would not change CPM prediction performance. However, if the generated connectome better predicts individual behavior compared to the empirical task-free connectome, this will indicate that the C2C models would extract a hidden reliable pattern of connectome reorganization from task-free state to task-specific state, to generate connectomes of task-specific states.

The behavioral predictive power of the model-generated task connectomes was significantly higher than the task-free connectomes. In WM task states, the predictive power of the model-generated connectome was r=0.180, and the power of the task-free connectome was r=0.076 *(p < 0.01)* (**Figure 4A, S6A, and S7**). This result held for all other task states *(all p’s < 0.01),* demonstrating that model-generated task-specific connectomes have stronger predictive power than the empirical task-free connectomes. Thus, in the absence of task-involved data, the C2C models can transform task-free data to provide more accurate behavioral predictions.

**Figure 4.**
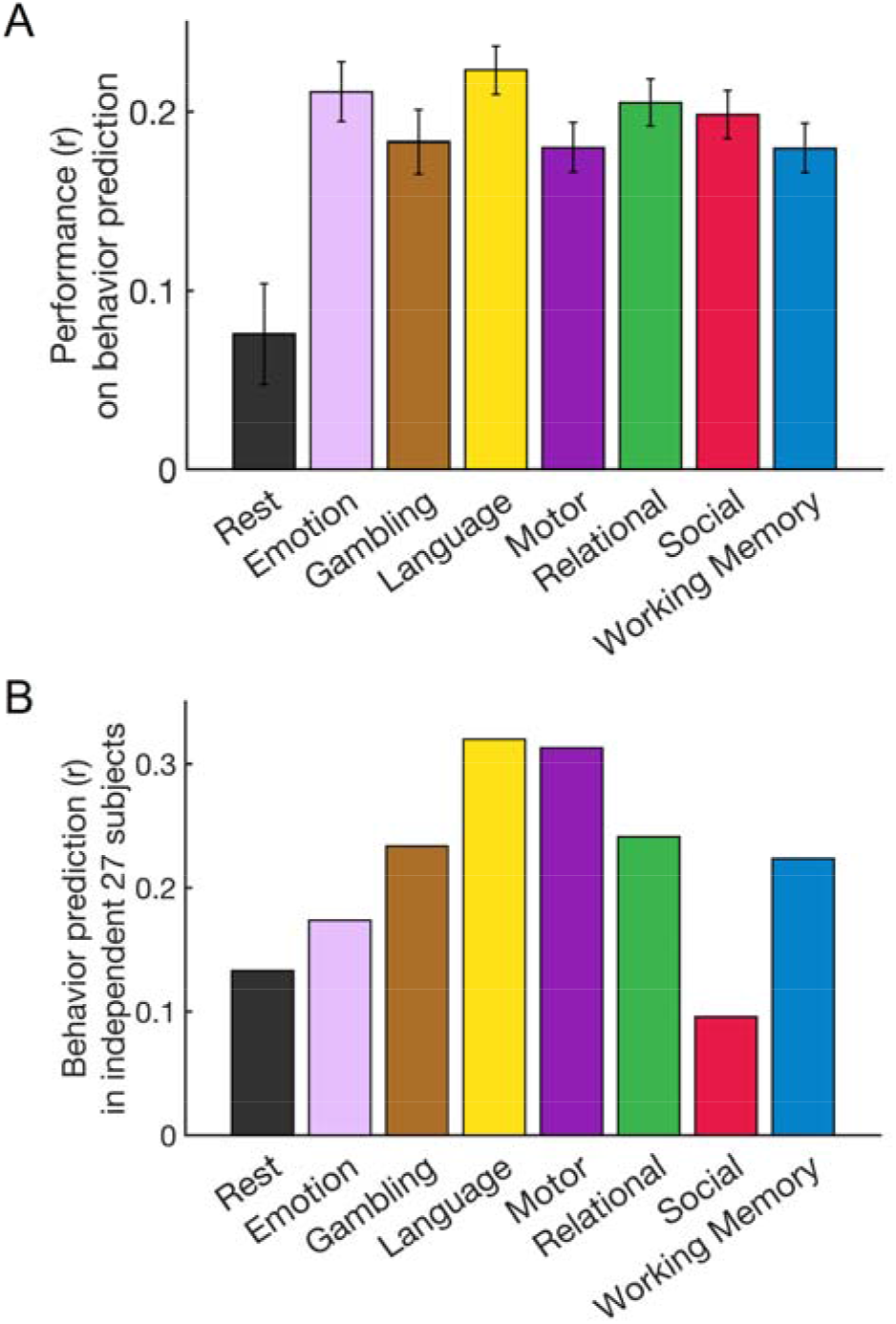
Individual behavior prediction with generated task connectomes. A) Predictive power for individual differences in fluid intelligence by the generated task connectomes (colored) in comparison to the observed task-free connectome (black). Error bar represents standard deviation from 1,000 iterations. The predictive power of the empirical task connectomes that can be considered practical ceilings is represented in **Figure S7**. B) Behavioral prediction with generated task connectomes of 27 subjects who did not have seven complete task scans. The left-most black bar shows the predictive power of the observed task-free connectome as a baseline. Prediction performance was assessed by correlating predicted scores with observed scores.

We also directly compared the similarity between the model-generated and the empirical task connectomes within individuals relative to across individuals. We performed this comparison for all task states and found that within-individual similarity is higher than cross-individual similarity for every task (**Figure S8**). Higher within-individual similarity and better intelligence prediction suggests that the C2C models generate task connectomes in a way that not only preserves but amplifies individual uniqueness.

### C2C model can generate task-specific connectome of individuals who have never undergone task fMRI

We performed additional validation of the C2C model in 27 subjects who were excluded in the main analysis because they were missing some task scans. Since this sample did not have the empirical task fMRI, we could not assess the accuracy of C2C models’ generating individual connectomes. It was only possible to compare the behavior predictive power of the model-generated connectome with that of the empirical task-free connectome. We did replicate the CPM result in this sample by revealing that, compared to the task-free connectome, the model-generated task connectomes provided better predictions of individual fluid intelligence scores (**Figure 4B and S6B**).

### Understanding the large-scale brain reorganization between cognitive states in a qualitative and quantitative way

We next sought to understand the systematic reorganization of the whole brain connectome across cognitive states using the connectome transformation framework. The constructed and validated C2C models provide a window that allows us to quantitatively characterize connectome reorganization between cognitive states. We hoped to provide a way to expand our understanding on the functional mechanism of the large-scale brain network in supporting diverse human cognition. We first compared subsystems (here, PCA components) of the whole-brain connectome between cognitive states, and then investigated composition of task-specific subsystems relative to task-free state subsystems.

**Figure 5 and S9** visualize the first 25 principal components of the task-free state. Here, all 268 brain nodes were divided into eight canonical networks (1: medial frontal, 2: frontoparietal, 3: default mode, 4: subcortical cerebellum, 5: motor, 6: primary visual, 7: secondary visual, 8: visual association networks, from Finn et al., 2015). The first component corresponds to the group mean of the whole-brain connectome. This component principally defines within-network connectivity of eight networks, consistent with previous observations, such as the resting-state networks extracted by independent component analysis on fMRI time-series or the modular structure revealed by graph theoretical approaches on the functional connectome. The following components revealed more distributed connectivity across networks, primarily defining cross-network connectivity. We compared all task-free components with task-specific components of each task state. In doing so, we sorted the order of task-specific components to be consistent with the task-free components. In other words, reordered *i*-th task-specific component has the spatial distribution maximally similar to the *i*-th task-free component, establishing a correspondence (ideally, one-to-one) of components between cognitive states. We are aware of that multiple task-free components could be combined to produce one task-specific component or that one task-free component could be divided into multiple components in task-specific states. We, however, stuck to building this correspondence for simplicity in identifying and comparing subsystems from different cognitive states.

**Figure 5.**
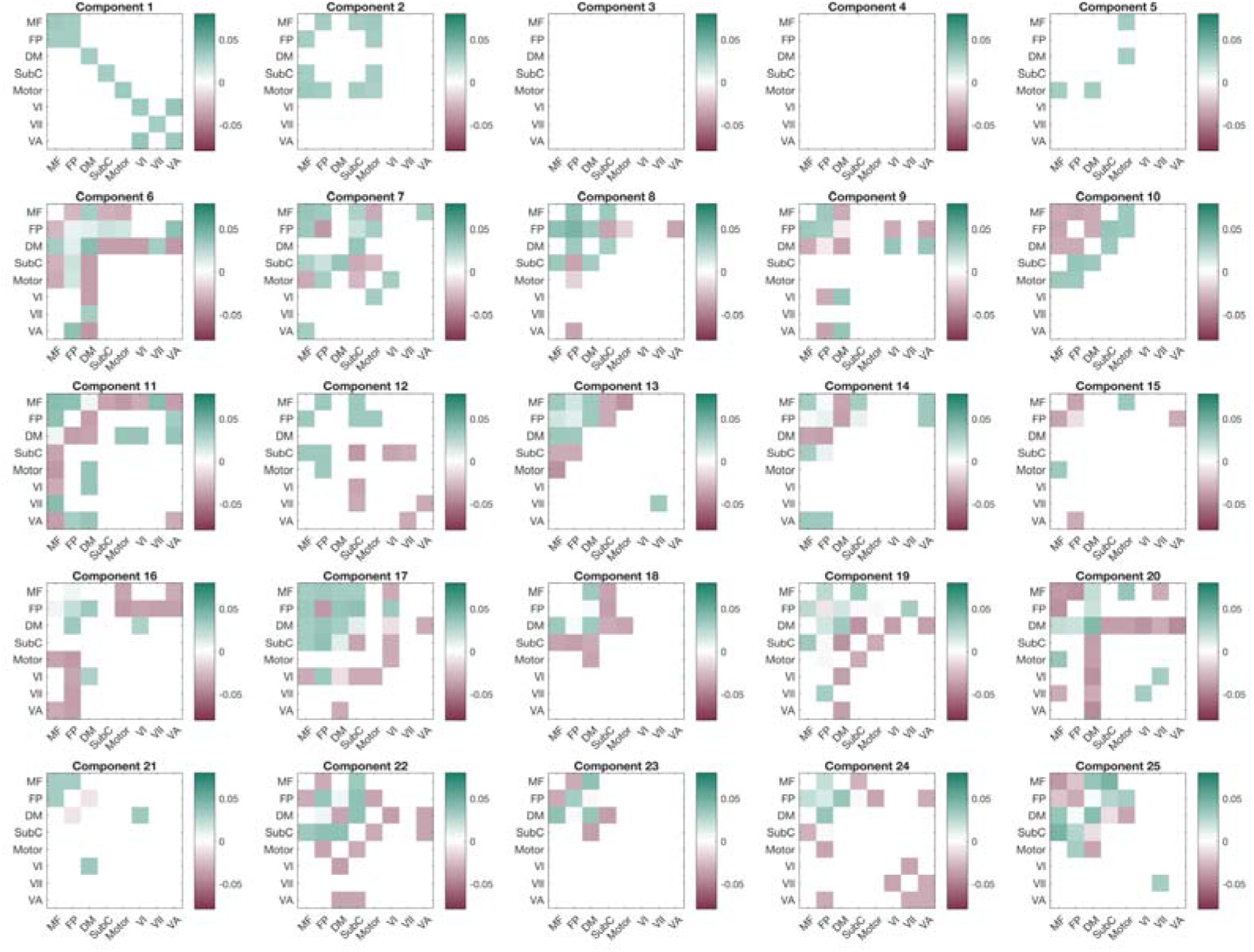
The first 25 task-free components from PCA. All components were thresholded *(Z* > 5.7\**SD*) at the edge-level first and then presented with eight canonical networks. The first component corresponds to the group mean of the whole brain connectome, and principally defines within-network connectivity of eight networks. The following components revealed more distributed connectivity across networks, primarily defining cross-network connectivity. Blank (by white color) indicates that all edges between two networks have subthreshold PCA weights.

### Task-general components relative to task-free state

**Figure S10** presents the similarity of the first ten components from each state with the first ten task-free components. Across all seven task-specific states, the first two components have apparently high similarity to the corresponding task-free components. The first component represents group-common state-general components (the group-mean structure as described earlier). It should be noted that the fact that a component is common to group does not necessarily imply that its weight is the same across subjects. Individual subjects could still have varying weights for group-common components.

We sought to quantify the large-scale reorganization with cognitive states. In the C2C state transformation, every task-specific component is described by combination of task-free components and the combination is defined and represented by PLS coefficients in C2C modeling. Hence, we compared the coefficients of the *i*-th component between seven task-specific states, for all *i* from *1* to *100,* to uncover task-general and task-specific reorganization (**Figure 6**). As can be expected, the first component of each task-specific states has distinctively strongest weights on the first task-free component. In contrast, this component has relatively negligible weights on the other task-free components. This pattern of coefficients distribution indicates that the first component is preserved in cognitive states transformation from task-free state, and thereby can be considered a state-general component across all task-specific states as well as task-free state.

**Figure 6.**
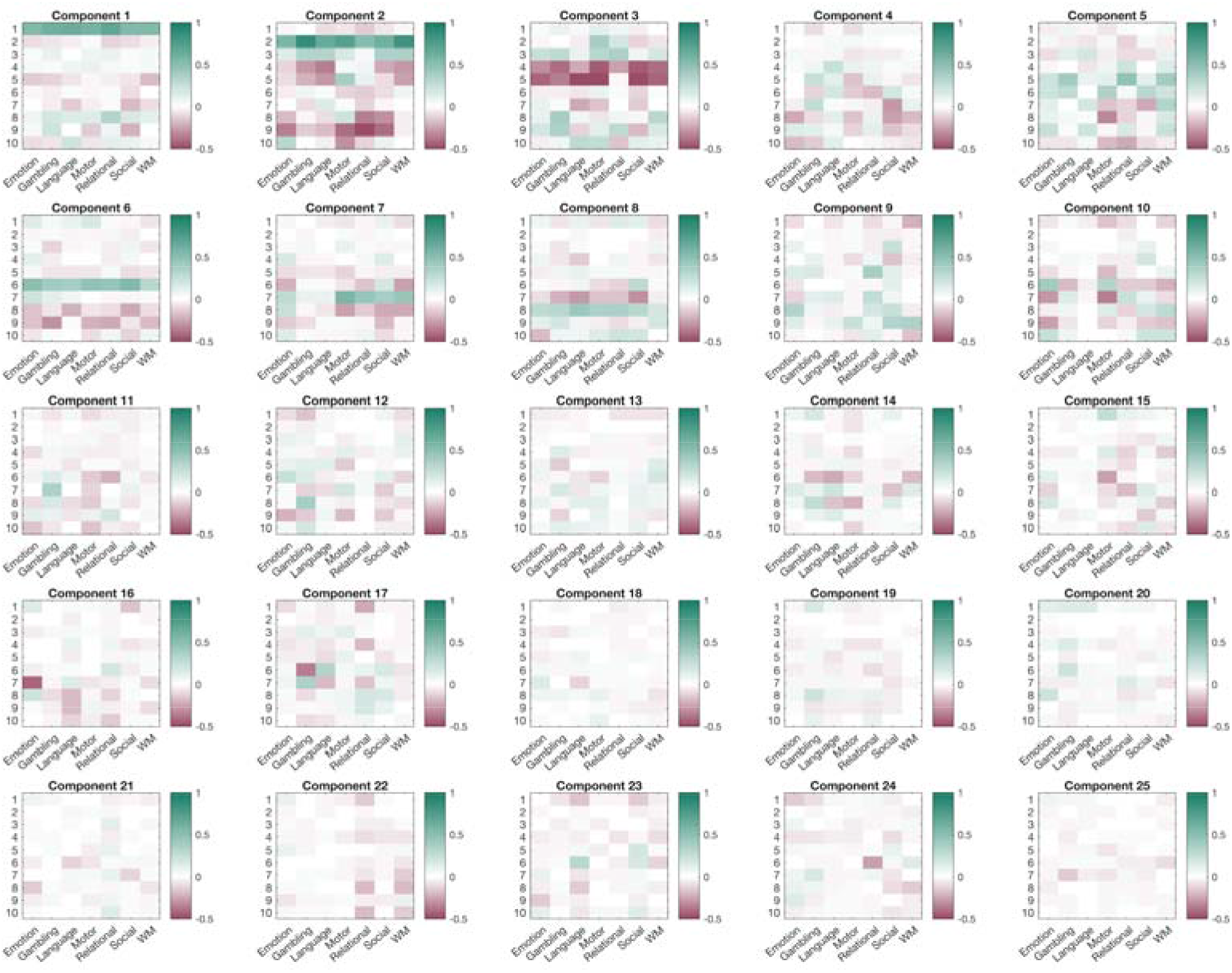
PLS coefficients presenting reorganization from task-free components (1-10 in each cell) to task-specific components (1-25, plotted across the figure). These coefficients describe composition of task-specific components. For example, the first task-specific component has the strongest weight on the first task-free component, whereas they have relatively negligible weights on the other task-free components. This suggests the first task-specific components as state-general components across tasks and task-free states. The second task-specific component has similarly the strongest weight on the corresponding second task-free component, however it also has relatively strong weight on another task-free components. For example, the second component of relational and social task states has strong negative weights on the eighth and ninth task-free component. This suggests that the second task-specific components are more or less state-general on a continuum, but not as much as the first component. In this figure, coefficients for only the first ten task-free components were visualized.

Importantly, we also found that there are other components (for instance, component 2, 3, 6, 7, and 8) that change consistently across tasks but do not solely correspond to a single task-free component (**Figure 7A**). That being said, these components are similar across task-specific states and exhibit the domain-general difference relative to the corresponding task-free state. These components had significantly similar distribution of weights on task-free components (**Figure 6**). Noteworthy, component 6 primarily represents default mode network’s within-network connectivity as well as connectivity with other networks (**Figure 5**). The component 6 also defines medial-frontal and frontoparietal connectivity in a moderate degree. The component 6 has significantly similar beta coefficient between almost every pair of seven cognitive states. While this component has the strongest weight on the 6-th component of the task-free state for all seven states, it also has comparable degree of weights on different components (for instance, task-free component 8 and 9 with the opposite direction **in Figure 6**). Other components (2, 3, 7 and 8) also exhibited relative task generality, although there are not fully general across all tasks (**Figure 7AB**). Among these networks, component 2 represents connectivity between medial frontal and motor networks as well as their connectivity to frontoparietal and subcortical-cerebellum networks. In addition, component 7 mainly represents the subcortical-cerebellum connectivity, and component 8 has the frontoparietal connectivity and, in a lesser degree, subcortical-cerebellum connectivity.

**Figure 7.**
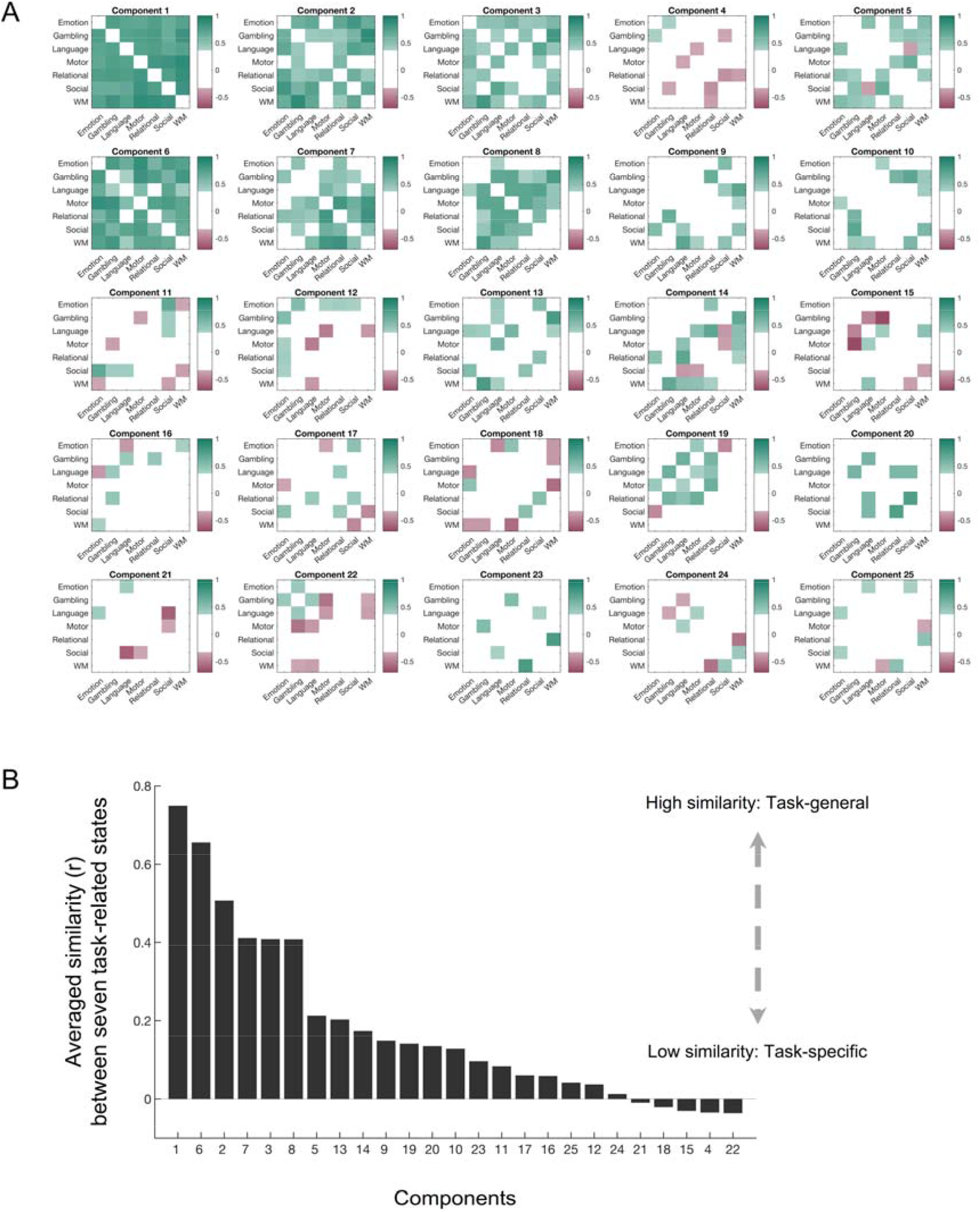
A) PLS coefficient similarity of components between task-specific states (threshold p < 0.001). B) Averaged similarity between seven task-specific states. Higher similarity indicates that a component is task-general, and lower similarity suggests that a component is task-specific on a continuum.

### The functional brain connectome exhibits task-specific components as well

Although the components described above showed task generality, the other components (4, 5, 9 and later) exhibited task specificity. These components had low similarity (*r* ~< 0.2) between task-specific states in their coefficients on task-free components (**Figure 7AB**). Among them, component 9, for instance, represents connectivity of frontoparietal network and connectivity of the default mode network, and component 11 primarily represents connectivity of medial frontal network (**Figure 5**). These components also did not have a distinctive weight on any task-free component, suggesting that these components are appeared from a distinct combination of multiple task-free components (**Figure 6**). This observation suggests that some functional components, especially connectivity of frontal and parietal brain areas, change in different ways across different cognitive states; in other words, these components are task-specific.

Note that brain networks or regions can be involved in multiple components. What defines a component is not the participant networks themselves, but rather the pattern of interaction between networks. For example, frontoparietal and default mode networks seem to play the main role in components 6 and 9, which were considered task-general and task-specific, respectively. The difference is the pattern of their connectivity with other networks. Only component 9 consisted of the opposite pattern of connectivity between the two networks. Moreover, component 6 additionally included the connectivity of the medial-frontal network.

## Discussion

Here, we presented a new connectome-to-connectome (C2C) state transformation modeling approach that generates individual task-specific connectomes. Rather than relying on regional response patterns to cognitive demands, the proposed model generates inter-regional interaction patterns, the functional connectome, for each subject across a wide range of cognitive domains, solely from the task-free connectome. The C2C state transformation model demonstrates both a high degree of *task specificity* across seven task states and *individualization* that closely resembles empirical task connectomes across individuals. Moreover, the model amplifies behaviorally relevant individual differences in task-free connectivity patterns, thereby improving prediction of individual differences in cognition such as intelligence.

The principal aim of this study was to develop a computational model to understand the connectome reorganization between cognitive states at the individual level. As a result, the C2C modeling in this study consists of relatively simple linear functions, principal component analysis and partial least square regression, for estimating connectome reorganization. Recent studies have revealed the high degree of functional connectome similarity between different cognitive states, integrative hubs playing a role across cognitive domains as well as task-specific alterations of functional connectome (Cole et al., 2013, 2014; Gonzalez-Castillo et al., 2015; Gratton et al., 2016, 2018; Shine et al., 2016; Shirer et al., 2012). These studies have contributed to the literature in a meaningful way that describes the large-scale commonalities and differences in functional connectome between various cognitive states. However, the field lacks a principled approach to characterizing task-induced modulation of functional connectivity. To address this, we designed a C2C model that can generate individual state-specific connectomes for seven different cognitive states. Furthermore, this model is transparent and interpretable, so that we can scrutinize model-estimated reorganization to infer brain reorganization.

In this study, we defined task-specific functional connectivity as synchronized fluctuations between brain regions during task performance. The task-specific functional connectivity, however, could be further disassembled into task-induced “true” modulation in connectivity, task-independent connectivity on which task engagement induces modulation, and event-related simultaneous activations of brain regions. There would exist “intrinsic functional connectivity” that closely resembles task-free connectivity and engaging in a cognitive task alters the intrinsic connectivity, yielding so-called extrinsic or task-evoked connectivity (Cole et al., 2014). Furthermore, task coactivations induces spurious, but systematic inflation in connectivity (Cole et al., 2019). It is important for C2C modeling to generate task-evoked connectivity, not task-related coactivation between brain regions. Task coactivation might largely relate to the experimental task design to which brain regions response in common, not the cognitive processes required in the task. In this study, we demonstrated that the C2C model-generated task-specific connectomes better predict individual intelligence compared to the task-free data. Our observation of improved behavioral prediction indicates that the C2C modeling actually predicts task-evoked connectivity. Greene et al., 2020 showed that task coactivation cannot predict intelligence, whereas task-dependent and task-independent connectome contribute to accurate prediction (Greene et al., 2020). That being said, if the C2C model only predicted the pattern of task-coactivation on the top of the intrinsic connectivity, then the model-generated connectome would have predicted individual behaviors with accuracy similar to the empirical task-free connectome or lower. This means that the presented C2C models truly predict (at least a portion of) task-induced modulation in the connectome. It should be noted, however, that these observations do not necessarily indicate that the C2C model generates task-evoked connectivity exclusively, as task-related connectomes generated from C2C state transformations would contain the effect of task coactivation as well.

Predicting task connectomes from task-free connectomes should have important implications for basic and clinical research. First, our C2C transformation models can inform how the brain reconfigures to support our cognitive and mental functions across tasks. Specifically, our method can help dissociate domain-general networks that play a role in diverse tasks from task-specific networks that are exclusively involved in different tasks. Second, the C2C approach takes advantages of task-free and task data, potentially providing a practical benefit. Task-free scans are easier to collect consistently across studies and sites than task scans. For example, patient groups or populations may have difficulty performing certain tasks (Pujol et al., 1998). Instead, task-free scans can be acquired because of their simplicity and minimal demands (Bullmore, 2012). Although task-free scanning offers the ease of acquisition, it is limited for characterizing individual traits and behaviors because participants can engage in unconstrained, subjective mind-wandering during scanning, making mental states more variable from scan to scan. Higher consistency across subjects and sessions can be obtained by requiring subjects to perform a common, explicit task, or to watch naturalistic movies (Finn et al., 2017; Vanderwal et al., 2017). In a task-engaged setting, participants are supposed to employ the same, or at least similar, cognitive functions to achieve a common task goal. Thus, task-induced consistency in the brain would help to better construct a brain-behavior association (Greene et al., 2018; Yoo et al., 2018). Through the additional validation, we tested the constructed C2C models in the novel set of subjects who have not completed task scanning. In these samples, the model-generated task-specific connectomes outperformed the empirical task-free connectome in predicting individual intelligence. This result shows that the proposed C2C modeling presents the strengths of task-free and task-specific data.

We note that while the current study contributes to the field with a keystone to study connectome reorganization, it needs further elaboration in future studies in several respects. First, the currently demonstrated modeling starts from several assumptions, such as one-to-one correspondence between cognitive states and the constant number of subsystems across states. While these assumptions help simplifying model construction as well as interpretation, they may differ from the way in which the human brain reorganizes. The brain network exhibits a hierarchical structure and modular organization (Meunier et al., 2010; Sporns and Betzel, 2016). Moreover, multiple subnetworks could be dynamically combined into one larger subnetwork, or vice versa, during cognitive engagement (Shine et al., 2016, 2019). The currently proposed model does not take account into such a network integration and segregation. Procedures defining subsystems or their relation between states could be overlooked as parameter optimization in regard to machine learning. We should, however, keep in mind that this framework is to model the human brain, not to build the best working machine. In this sense, it could deepen our understanding of the cognitive brain by incorporating biologically-driven settings and constraints into state transformation modeling.

## Author contributions

Conceptualization, K.Y. and M.M.C., Methodology, K.Y., Software, K.Y., Validation, K.Y. and Y.H.K., Formal Analysis, K.Y., Y.H.K., M.D.R., and D.S., Writing – Original Draft, K.Y., M.M.C., M.D.R., and Y.H.K., Writing – Review and Editing, K.Y., M.D.R., M.M.C., D.S., and R.T.C., Visualization, K.Y., M.M.C., M.D.R., and D.S., Funding Acquisition, M.M.C.

## Acknowledgements

This project was supported by National Institutes of Health grant MH108591 and by National Science Foundation grant BCS1558497. Data were provided by the Human Connectome Project, WU-Minn Consortium (Principal Investigators: David Van Essen and Kamil Ugurbil; 1U54MH091657) funded by the 16 NIH Institutes and Centers that support the NIH Blueprint for Neuroscience Research; and by the McDonnell Center for Systems Neuroscience at Washington University.

## Materials and Methods

### MR data – Human Connectome Project S1200

We obtained minimally pre-processed MRI data from the S1200 release of the Human Connectome Project (HCP) (Essen et al., 2013; Glasser et al., 2013). This dataset contains nine fMRI conditions per subject including seven tasks (emotion, gambling, language, social, motor, working memory [WM], and relational) and two separate rest conditions during two-day visits. Each condition involves two runs with opposite phase encoding directions (LR and RL). All fMRI data were acquired on a 3 T Siemens Skyra using a slice-accelerated, multiband, gradient-echo, echo planar imaging (EPI) sequence (TR = 720 ms, TE = 33.1 ms, flip angle = 52°, resolution = 2.0mm^3^, multiband factor = 8). Detailed information on MR imaging parameters and preprocessing procedure have been published elsewhere (Barch et al., 2013; Smith et al., 2013; Uğurbil et al., 2013). The experimental protocol was approved by the Institutional Review Board at Washington University in St. Louis.

The main analysis in this study was limited to 316 participants. Out of 1,206 subjects, we first selected 561 individuals who completed all nine fMRI scans (888/1206), exhibited low head motion in all fMRI runs (<3mm translation, <3° rotation, and <0.15mm mean frame-to-frame displacement) (565/888), and had a behavioral fluid intelligence score (561/565). The final pool of subjects was composed of 316 subjects who were unrelated to any of the other 316 subjects based on their family structure verified with genetic information (316/561). In this control of relatedness, we randomly selected a single subject from each family and excluded all the other family members.

We performed additional preprocessing steps on the minimally pre-processed fMRI scans. The first 15 volumes in each run were discarded. Nuisance covariates were regressed from each run using custom scripts in MATLAB R2016b. Nuisance covariates included 24 motion-related parameters (6 translational and rotational motions, 6 derivatives, and their squares), three mean tissue signals (global, white matter and cerebrospinal fluid), and linear and quadratic trends.

### Construction of the Whole-Brain Functional Connectome

A set of brain nodes covering the whole brain was defined with a 268-parcel functional atlas (Shen et al., 2013). An average time-series was extracted for each node from the preprocessed data. Pearson’s correlation between the mean time-series of every pair of 268 nodes was calculated as functional connectivity, providing a 268-by-268 whole-brain connectome. The same procedure was completed for every run and every state (i.e., task). The final rest connectome was constructed by averaging four rest-run connectomes (2 runs x 2 sessions). The final task connectome was constructed by averaging two task-run connectomes for each task.

### Connectome-To-Connectome (C2C) State Transformation Modeling

The presented C2C state transformation modeling is a generative model that can generate task-specific connectomes of individuals. In the current modeling, generation of task-specific connectomes is based solely on the whole-brain task-free connectome of individuals. Hence, this procedure provides a state transformation of the brain functional connectome.

A strength of the C2C state transformation model is that it is simple and transparent, which in turn, makes the model interpretable (Bzdok and Ioannidis, 2019). We sought to have this model as simple and interpretable as possible so that we can understand the large-scale mechanism and anatomy of the task transformations. The C2C model works in three steps. The first step is to extract task-free subsystems from the whole-brain task-free connectome of individuals. The second step is to transform task-free subsystems to estimate task-specific subsystems. The third step is to construct whole-brain task-specific connectomes. Importantly, C2C modeling generates the whole-brain connectome of targeted cognitive states at a single subject level.

### Model Construction

We constructed seven C2C models for all seven task states included in the HCP dataset. We constructed and validated models using 10-fold cross validation. In particular, we constructed C2C models using a training set (90%) of all available subjects and left out 10% as a testing set in which we validated the models. Detailed information on the 10-fold approach is provided in a designated section *10-fold Cross Validation.*

The proposed C2C state transformation model was constructed using two statistical methods, principal component analysis (PCA) and partial least square (PLS) regression. PCA was first employed to define and extract state-specific subsystems and their scores for the task-free state and task-evoked state, separately. In constructing the task-free to working memory (WM) task state transformation, for example, we performed one PCA using the task-free connectomes of individuals in the training set. This corresponds to the first step of the C2C model described in the previous section. We also performed another PCA separately using these same individuals’ WM task connectomes. This second PCA provides a reconstruction the whole-brain task connectome from the generated task subsystems, the third step of the C2C model. Then, PLS regression was employed to estimate the transformation of subsystems from the task-free state to the WM task state. The PCA-extracted subsystem scores of task-free and WM task states were put into PLS regression. We constructed one C2C model for each task state of the HCP dataset, producing a total of seven C2C task models.

To apply the C2C model to new task-free connectome data from individuals in the test set, as a first step, the model extracts individual scores of task-free subsystems that were predefined by the first PCA in model construction. In the second step, the model estimates individual scores of task-specific subsystems from task-free scores using pretrained PLS regression. Then finally, the model constructs the whole-brain task-specific connectome using the estimated task-specific scores and reconstruction that was also predefined by the second PCA in model construction.

The number of components in PCA was set to 100 based on previous studies reporting stable reconstruction of individual connectomes and accurate behavior prediction from between 50 to 150 components with a peak around 80 to 100 components (Amico and Goñi, 2018; Sripada et al., 2019). The C2C models with different numbers of components (50~200) produced similar results in this study.

### 10-Fold Cross Validation

We validated the C2C state transformation models using 10-fold cross validation. The C2C models were trained using nine folds of data (284 or 285 subjects) and tested on the one left-out fold (out-of-sample validation). Once we constructed the seven C2C models for the seven task states, we applied these C2C models to the task-free connectome in the left-out fold (31 or 32 subjects), generating seven (transformed) task connectomes, one from each model. Each of the 10 folds was iteratively left out as a test set in 10-fold cross-validation. We repeated this 10-fold cross-validation 1,000 times with randomizing subject-to-fold assignment to compute reliable statistics, demonstrating that the presented results are not dependent on specific data partitions. The models’ generated task connectomes were validated in multiple ways, described in the following sections.

### Similarity of Generated Task Connectomes to Empirical Task Connectomes

We first investigated whether the generated task connectome resembles the empirical task connectome more than the observed task-free connectome. To measure the similarity of the two connectomes, we computed the spatial correlation between them. The correlation describes the generative accuracy of spatial patterns at the connectome level. We additionally examine mean square error (MSE) between the generated and empirical task connectome. The MSE describes the generative accuracy of individual connectivity strength on average. The similarity was assessed in individuals and averaged across 316 individuals, and averaged across 1,000 iterations for each task.

### Task Specificity of Generated Task Connectomes

Next, we examined the task specificity of generated task connectomes in two ways. First, we compared the intra-state similarity between generated and empirical task connectomes with the inter-state similarity between them. Connectome similarity by spatial correlation was measured in individuals and averaged across 316 individuals, and averaged across 1,000 iterations of the cross-validation.

Secondly, we tested task specificity with a more conservative approach, connectome-based fingerprinting (Finn et al., 2015). Fingerprinting analysis aims to identify a participant from a group of individuals based on their unique functional connectivity pattern. We modified this approach in this study to test among states, not among individuals. State fingerprinting analysis requires two functional connectomes from each state: one (generated task connectome) to serve as the ‘target’ and the other (empirical task connectome) to serve as the ‘database’. Using the WM task as an example, the empirical WM connectome is considered successfully identified if the generated WM connectome maximally resembles the empirical WM connectome compared to the other six tasks’ empirical connectomes. In general, the empirical connectome of cognitive state *s* is considered successfully identified if

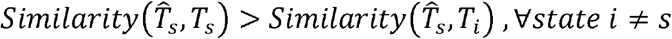

where 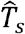 is the generated task connectome of state *s*, and *T_s_* is the empirical task connectome of state *s*. The similarity between two connectomes is measured by spatial correlation.

Then the measure of task specificity, identification success rate of a task, was measured as follows:

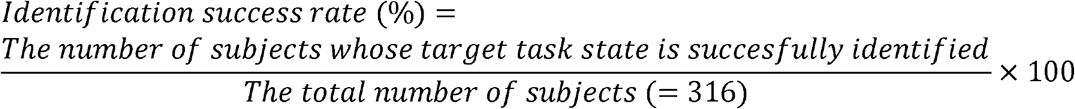

Success rate was computed for each task state in each iteration of 10-fold crossvalidation, and then averaged across 1,000 iterations to provide reliable statistics. A high identification success rate shows that the C2C modeling predicts task connectomes with a high degree of specificity.

We further tested if a high identification success rate of a certain task state is due to the high similarity of the empirical task-free connectome to the specific task connectome. To test this possibility, we related each individual’s task-free connectome to their task-state connectomes. Specifically, we computed the spatial correlation of the task-free connectome with each of the seven empirical task connectomes. Among the seven states, we tracked the task state of the connectome that was most similar to the empirical task connectome. After completing this analysis for all participants (*n*=316), we counted the number of participants for each state, providing a group-averaged similarity of task connectomes with the task-free connectome.

### Predictive Power of Generated Task Connectomes: Individual Difference in Fluid Intelligence

We next tested whether, compared to the empirical task-free connectome, the task connectomes generated by the C2C state transformation model had amplified behaviorally relevant information. We used data-driven connectome-based predictive modeling (CPM, Finn et al., 2015; Shen et al., 2017) to assess predictive power for individual fluid intelligence of the model-generated connectome relative to the empirical task-free connectome.

The CPM procedure was in principle the same as in our previous work (Yoo et al., 2018, 2019). The only difference from our previous studies is that here we trained the CPM model along with the C2C model. We trained a CPM on the same 10-fold crossvalidation loops in which the C2C state transformation models were trained. Thus, training samples, the empirical task and rest connectomes of the training set, for C2C modeling were also used to train the CPM. Once the two models were constructed, the C2C model first generated the task-specific connectome as described previously, and then the CPM predicted individual intelligence using the C2C-generated task connectome in the same left-out testing samples.

We assessed the predictive power of the generated task connectomes i) by correlating individual predicted intelligence scores with observed intelligence scores and ii) by generating prediction R^2^ based on mean square error (MSE, Scheinost et al., 2019). Behavioral predictions were also repeated with 1,000 iterations of 10-fold validation to provide reliable statistics. The predictive power of the task-free connectome was also assessed as a control.

### Additional Validation

We performed additional validation of the proposed C2C state transformation models in a set of HCP participants who were excluded in our main analysis because their task MR scans were incomplete. We again only used unrelated subjects (*n*=27) to rule out potential bias induced by family structure. In this validation, the C2C models for each task were trained with the entire set of 316 individuals used in our main analysis. The trained models were then applied to the task-free connectome of 27 new participants whose task connectomes were never analyzed in this study. The C2C models generated task-specific connectomes for each of these individuals. Since these 27 participants did not have complete task data, we could not assess the similarity or task specificity of generated task connectomes. We only examined whether the generated task connectomes better predicted individual intelligence compared to their task-free connectome.

### Subsystems of the whole-brain connectome

As described in *Model Construction,* PCA defines subsystems (i.e., principal components) in each task state. To compare the subsystems between different cognitive states, we first sorted the subsystems of task states to be consistent with the order of task-free subsystems. For simplicity, we assumed a one-to-one correspondence of subsystems between states. As an example for the WM task, we computed the spatial similarity of all WM task subsystems with the first task-free subsystem and then assigned the WM task subsystem that yielded the highest similarity to be the first subsystem of the WM task state. We next computed the spatial similarity of subsequent WM task subsystems with the second task-free subsystem and, similarly, assigned a WM task subsystem that yielded the highest similarity as the second. We repeated this procedure until all WM task subsystems were assigned. We ran the same sorting for all the other task states as well. This procedure allows to build, ideally, a one-to-one correspondence of subsystems between the eight task states (one task-free and seven task-specific).

### Understanding the Functional Reorganization of the Brain Connectome

We sought to reveal the relationship of task-specific states to task-free states by asking how task-free principal components reorganize into task-specific principal components. To study the reorganization patterns, we investigated the coefficients of PLS regression in the C2C models. Here, PLS coefficients represent a combination of task-free components in generating task components. Furthermore, they can be used to ask how state-specific vs. state-general the task-related components are. We first used a sorting procedure to match components across seven task states and rest, and then we compared the PLS coefficients of the components across these states. We assessed the similarity of coefficients between components by correlating their coefficients. High correlations indicate that the components of different states emerge from a similar reorganization of task-free components. In other words, high correlations demonstrate that, across different cognitive states, corresponding functional connectivity pattern principal components change from rest to task in similar ways. Along a continuum of similarity, high similarity across most pairs of task states would suggest that the subsystem is task-general, and low or variable similarity would suggest that a subsystem is task-specific.

## Data availability

The Human Connectome Project data used in the current study are publicly available at https://db.humanconnectome.org.

## Code availability

MATLAB script for the connectome-to-connectome (C2C) model construction is available to download at https://github.com/rayksyoo/C2C. Scripts for the other (statistical) analyses are available from the corresponding author upon request.

