## Supplemental Figures for "A cognitive state transformation model for task-general and task-specific subsystems of the brain connectome"

**Supplementary Figures**

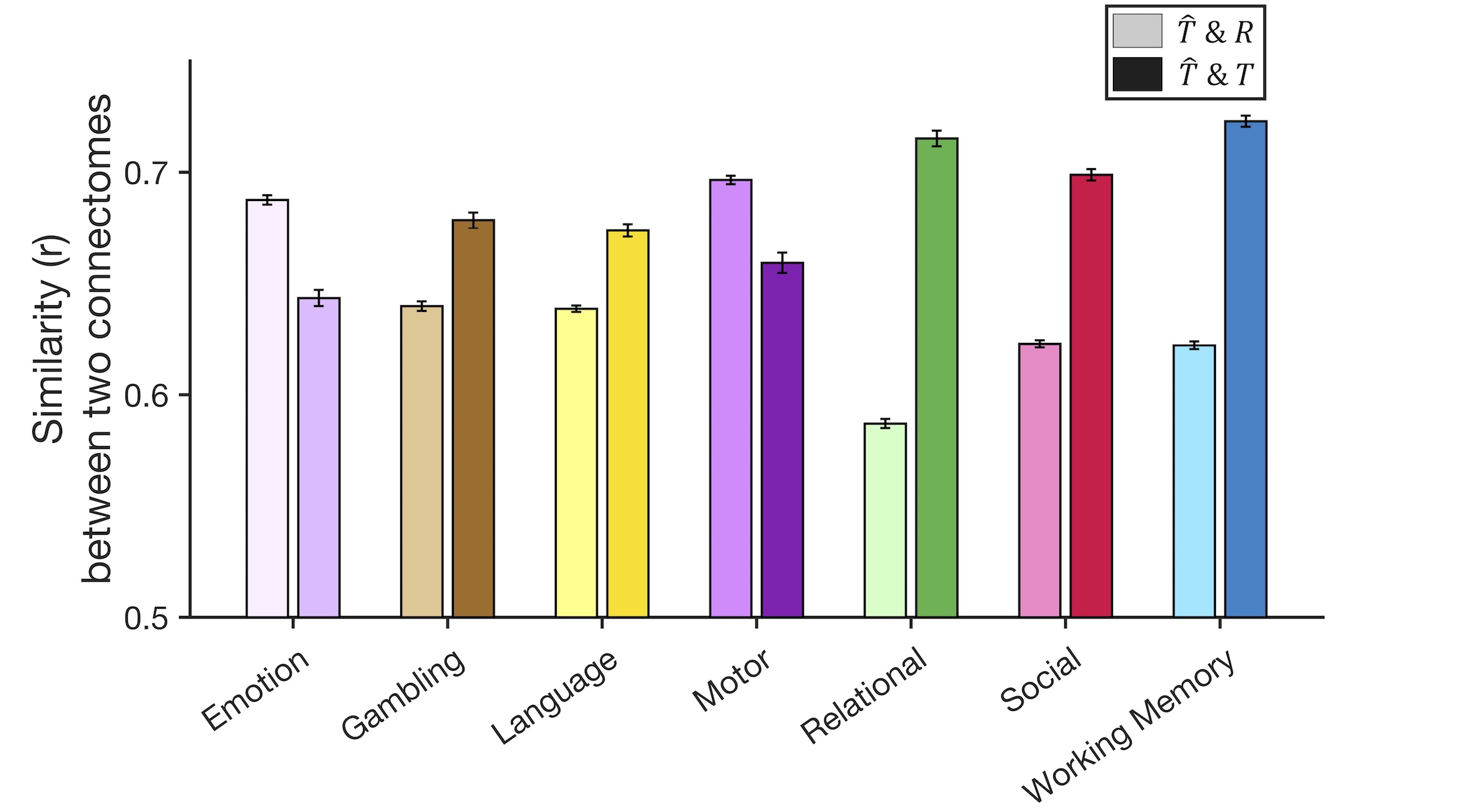

**Supplementary Figure S1**. Similarity of the generated task connectome to the observed task-free connectome and the observed task connectome. In each task column, a darker bar (right) represents a similarity between the generated and empirical task connectome, and a lighter bar (left) represents a similarity between the generated task and empirical task-free connectome. Error bars represent standard error across 316 subjects. ($R$: observed task-free (rest) connectome, $T$: empirical task connectome, $\hat{T}:$ generated task connectome).

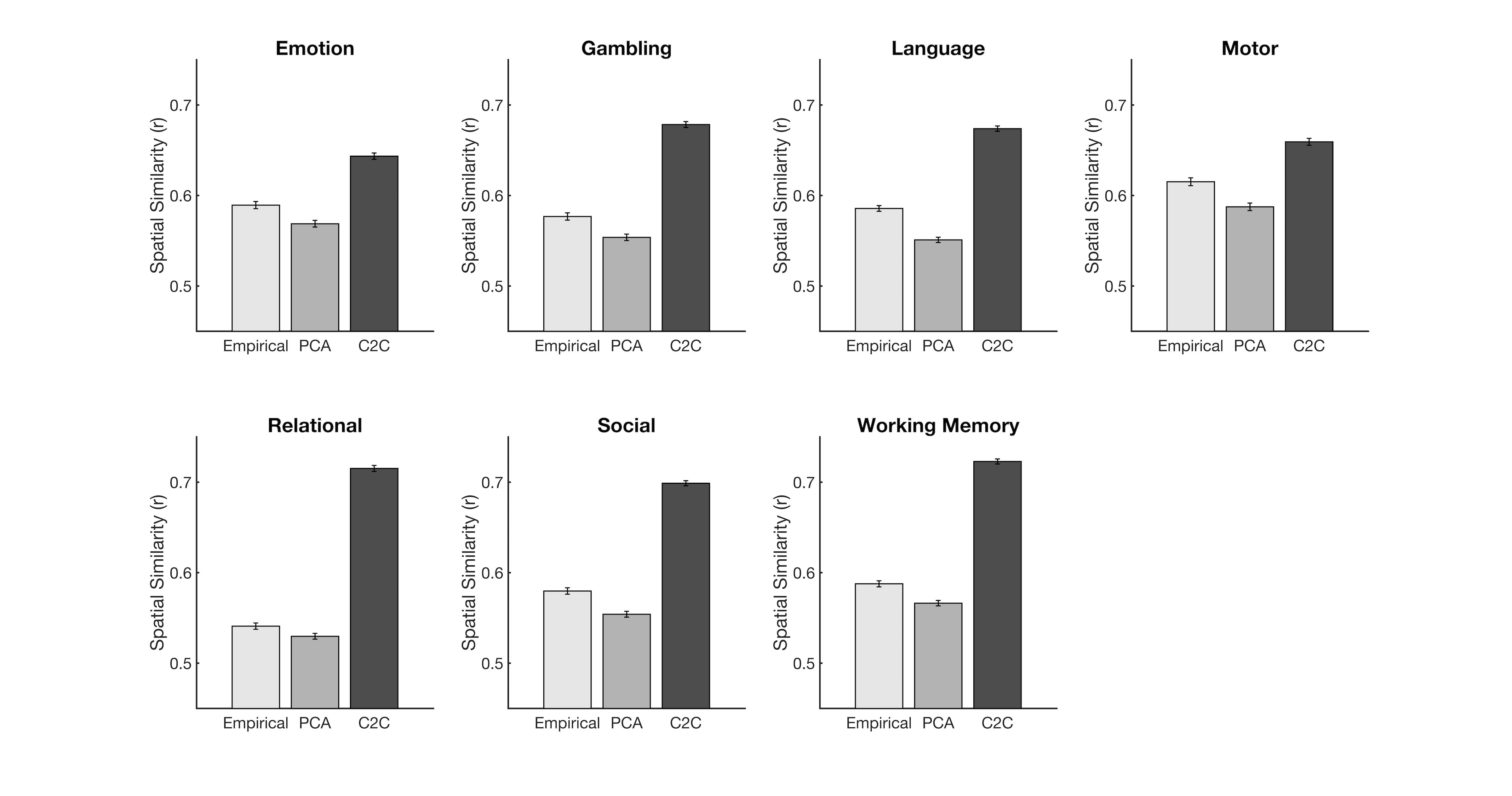

**Supplementary Figure S2.** PCA-based noise removal does not increase connectome similarity between task and rest states. This shows that the similarity between C2C-generated task connectome and empirical task connectome is statistically higher than that between the empirical (noise-removed) rest and task connectomes. Error-bars: standard error across 316 subjects.

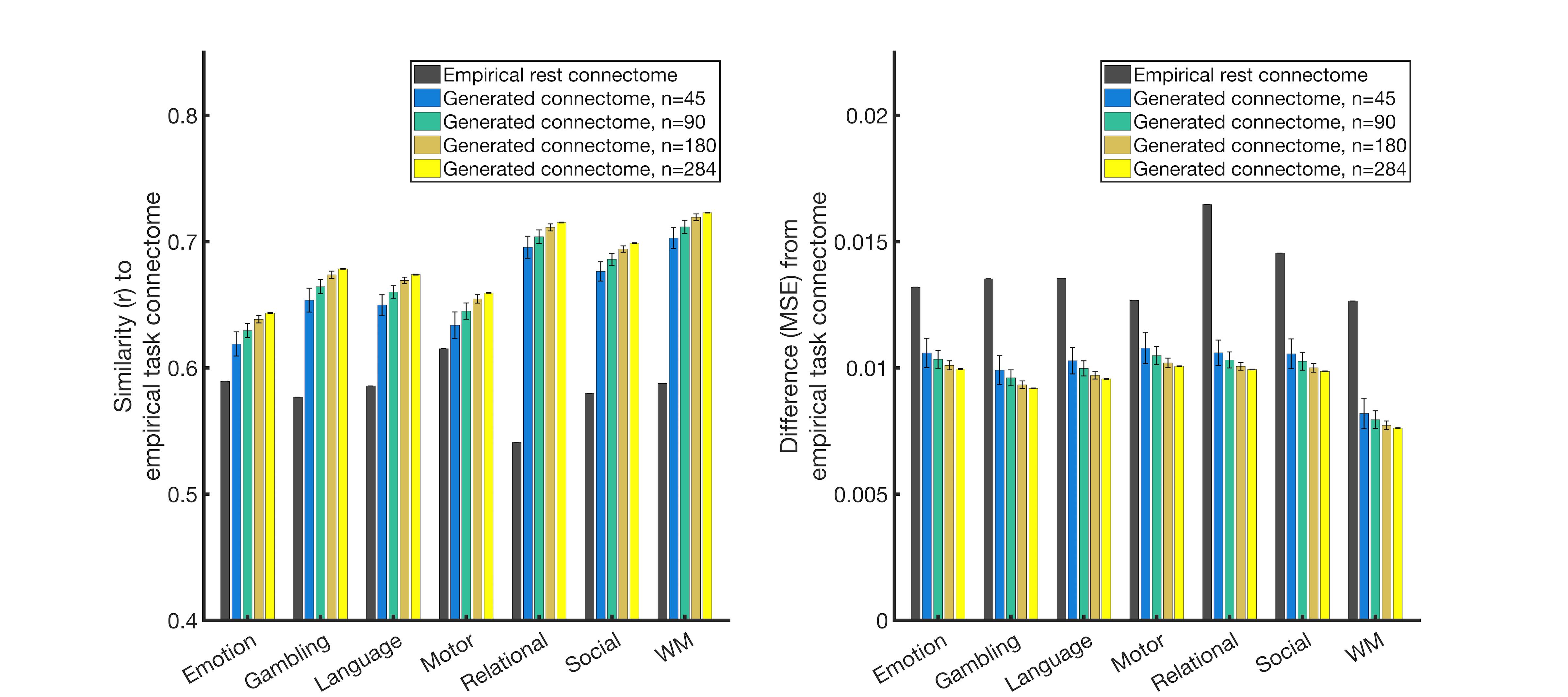

**Supplementary Figure S3.** Similarity of the C2C-generated task connectomes to the empirical task connectomes with different amount of training data. Increasing the number of subjects available to the C2C model training gradually increased model accuracy. Notably, even with a small number of training subjects (n=45), the C2C-generated task connectomes were significantly more similar than the rest connectome to the corresponding empirical connectomes. Error bars: standard deviation across 1,000 iterations.

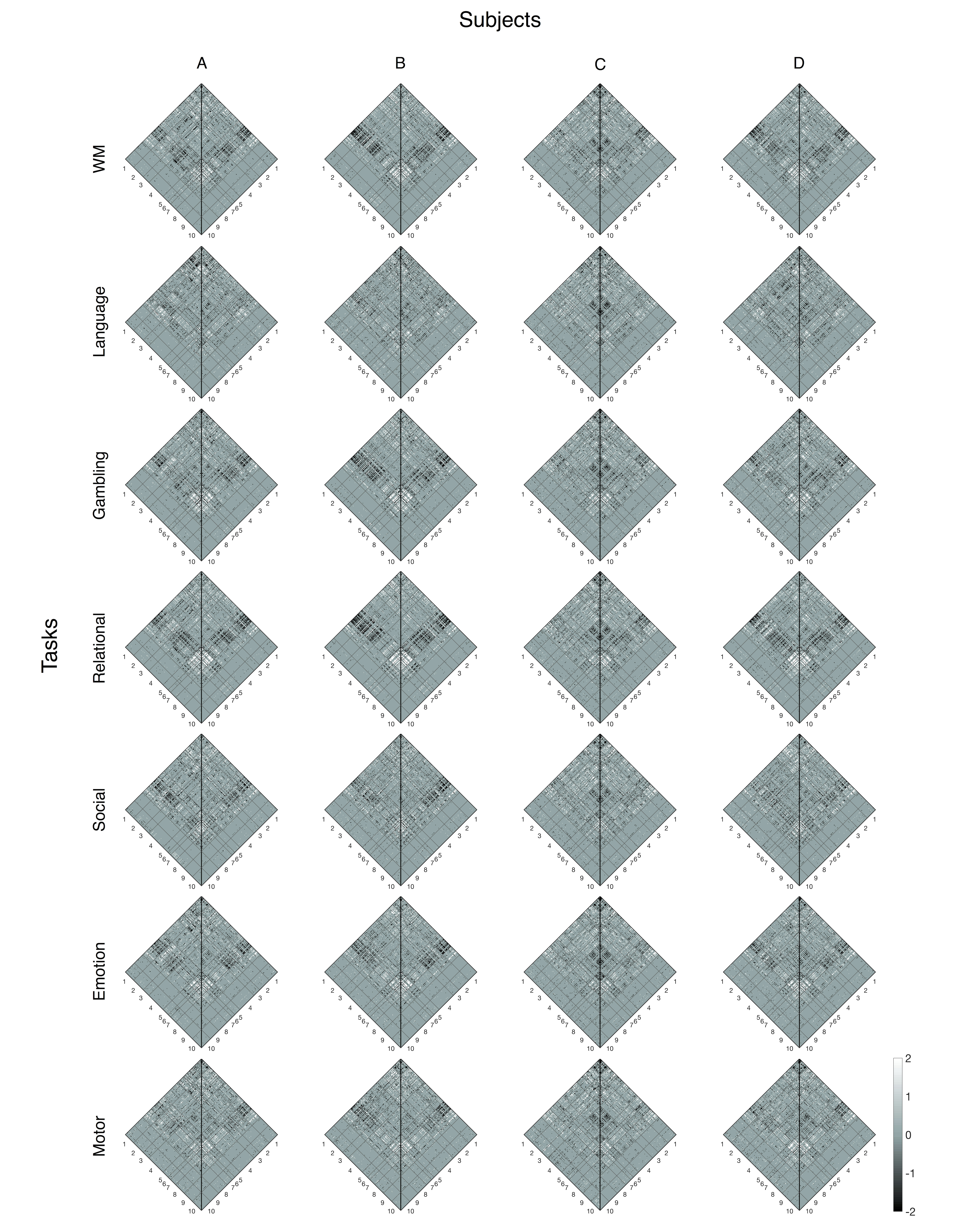

**Supplementary Figure S4. Visualization of the model-predicted difference and the observed difference between resting-state and each task state connectome from four selected individuals.** The C2C models accurately predict task-induced connectivity modulations across individuals and tasks. Columns correspond to four representative individuals and rows correspond to the seven tasks. In each connectome, the left triangle visualizes differences between the observed task connectivity and the observed rest connectivity (i.e. observed ‘task – rest’ contrast), and the right triangle visualizes differences between the C2C-generated task connectivity and the observed rest connectivity (i.e. predicted ‘task – rest’ contrast). The edge-wise differences were z-scored for each individual and state, and also thresholded at z=1.64 for visualization purposes. 1: Medial frontal network, 2: fronto-parietal network, 3: default mode network, 4: motor, 5: Visual I, 6: Visual lI, 7: Visual associations, 8: salience network, 9: subcortex, 10: cerebellum.

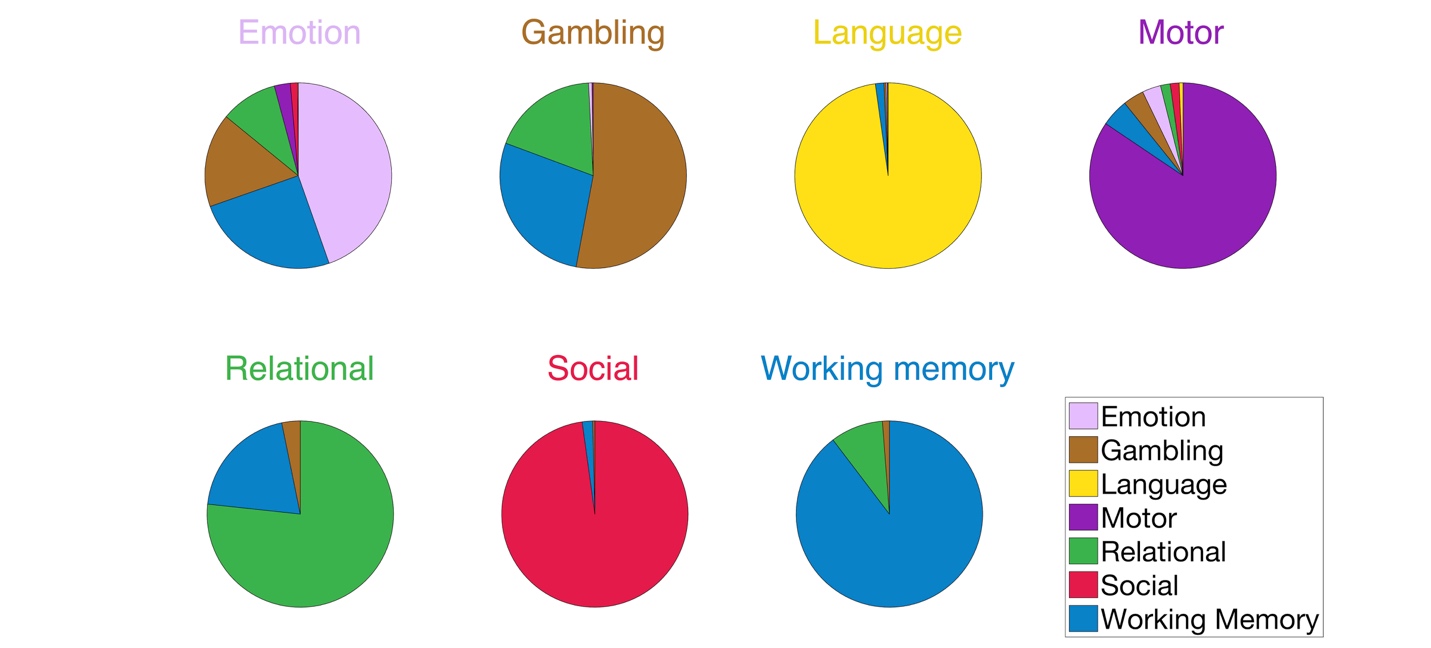

**Supplementary Figure S5**. Misidentification across seven tasks. Each color represents identification by each state. In every state, the largest part corresponds to the success identification of the empirical task connectome by the generated task connectome of the same task.

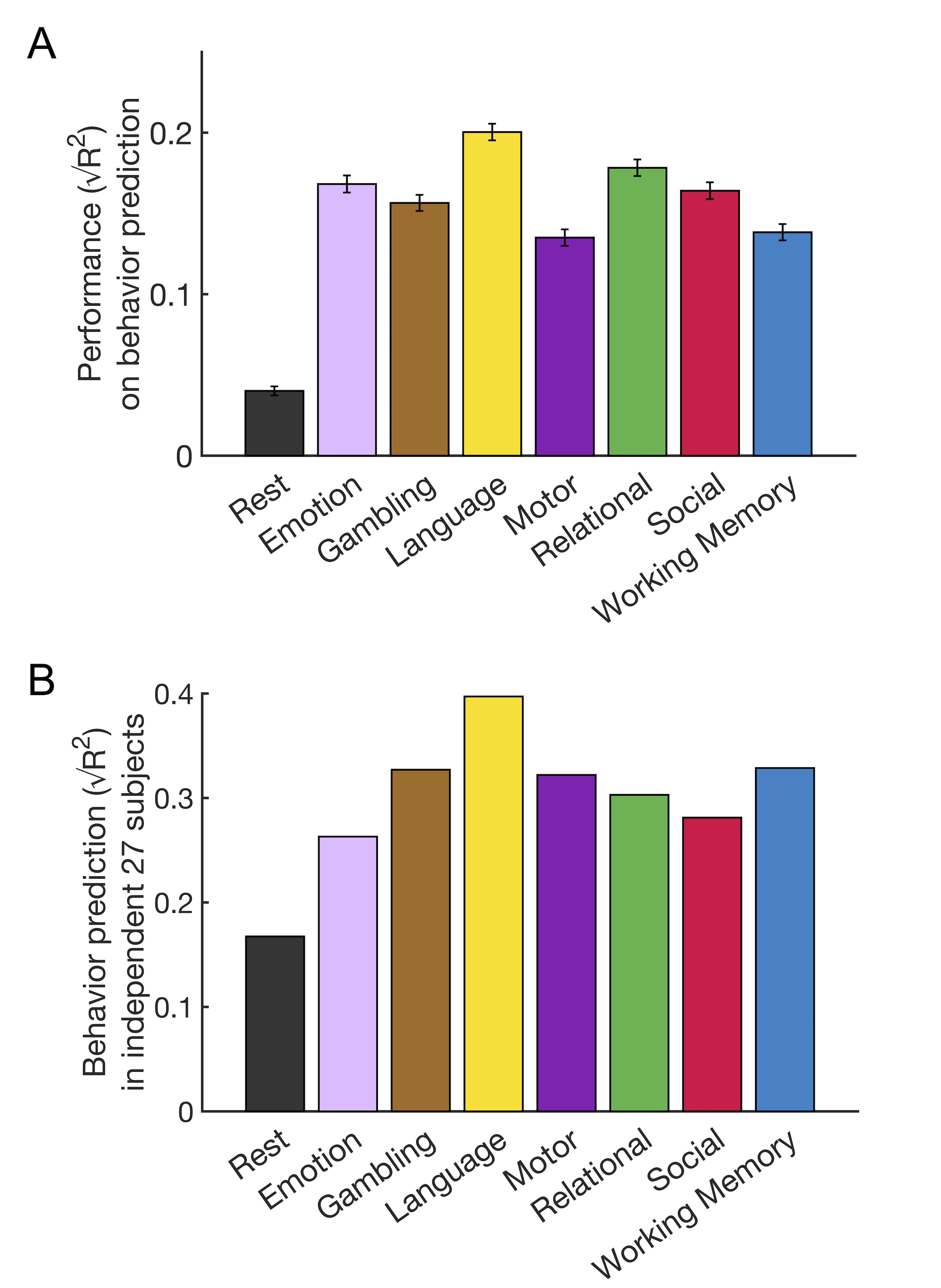

**Supplemental Figure S6**. Individual behavior prediction with generated task connectomes. A) This figure represents the same prediction as Figure 4A, but prediction performance was assessed by estimating prediction R^2^. Error bars represent standard deviation from 1,000 iterations. B) Behavioral prediction with generated task connectomes of 27 subjects who did not have seven complete task scans. The left-most black bar shows the predictive power of the observed task-free connectome as a baseline. This figure represents the same prediction result as Figure 4B, but prediction performance was assessed by estimating prediction R^2^.

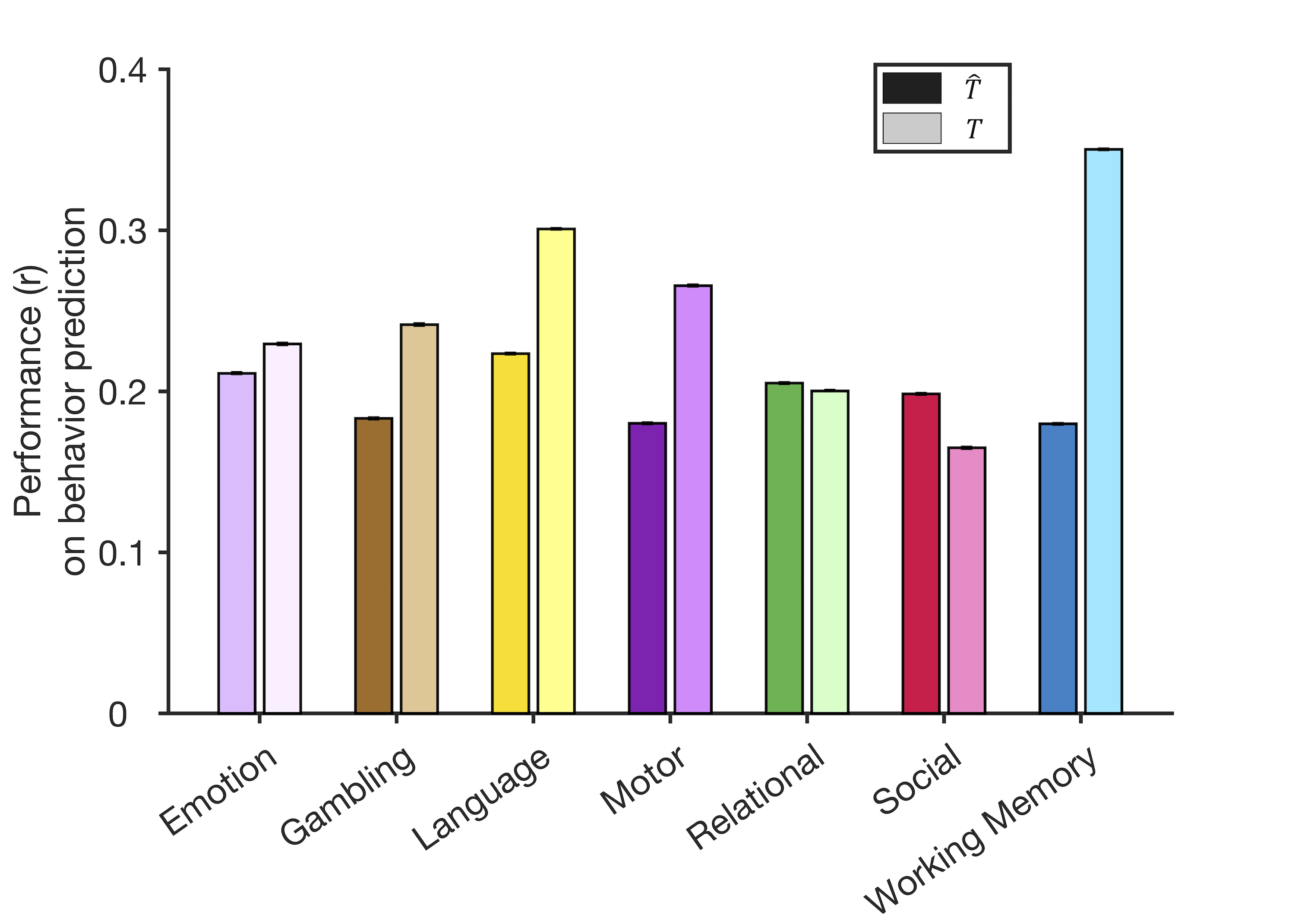

**Supplemental Figure S7**. Comparison of behavior prediction performance between generated task and empirical task connectomes. In each task column, a darker bar (left) represents the prediction performance of the generated task connectome, and a lighter bar (right) represents the performance of the empirical task connectome. Behavioral prediction with empirical task connectomes (lighter) represents theoretical upper limit of predictive power. Prediction performance was assessed by correlating predicted scores with observed scores. Error bars represent standard error from 1,000 iterations. ($T$: empirical task connectome, $\hat{T}:$ generated task connectome).

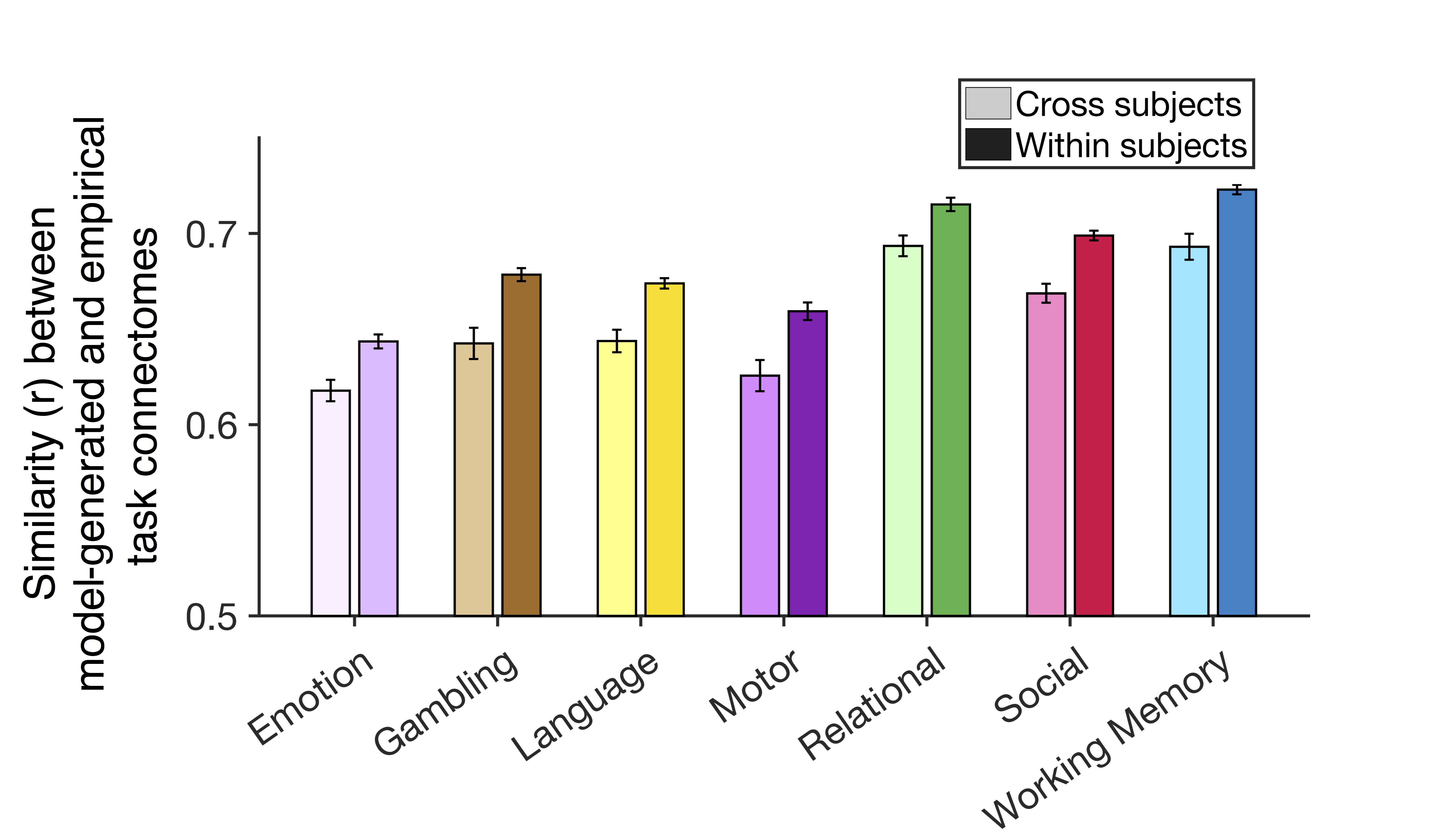

**Supplemental Figure S8**. Similarity between generated task and empirical task connectomes was higher in within-subject pairs than in across-subject pairs. In each task column, a darker bar (right) represents a similarity between the generated and empirical task connectome of the same individual, and a lighter bar (left) represents a similarity between the generated connectome of an individual and empirical connectome of a different individual. Error bars represent standard error across 316 participants.

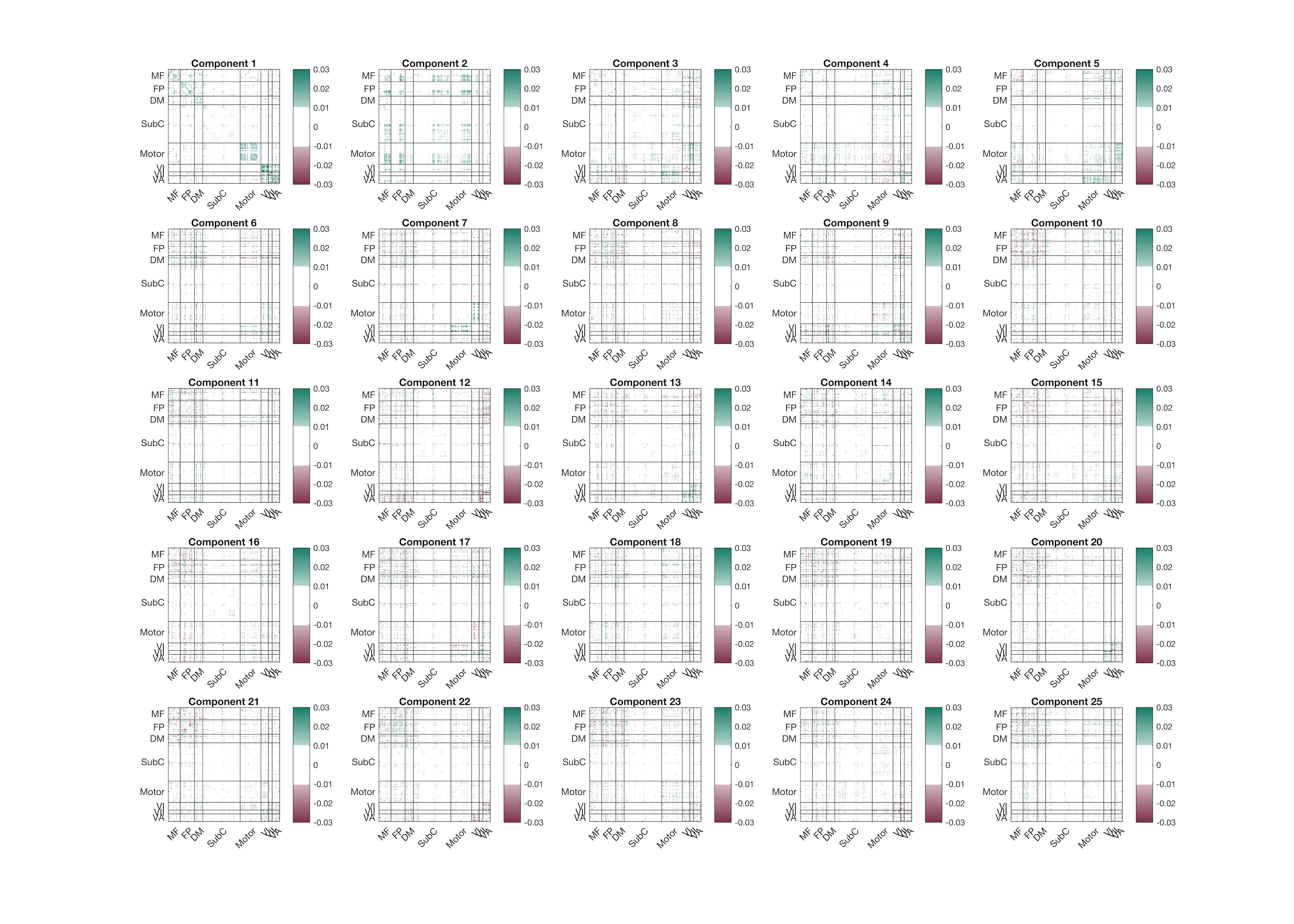

**Supplementary Figure S9**. The first 25 components of task-free connectome. The whole-brain components were thresholded (*absolute of a coefficient* > 0.01).

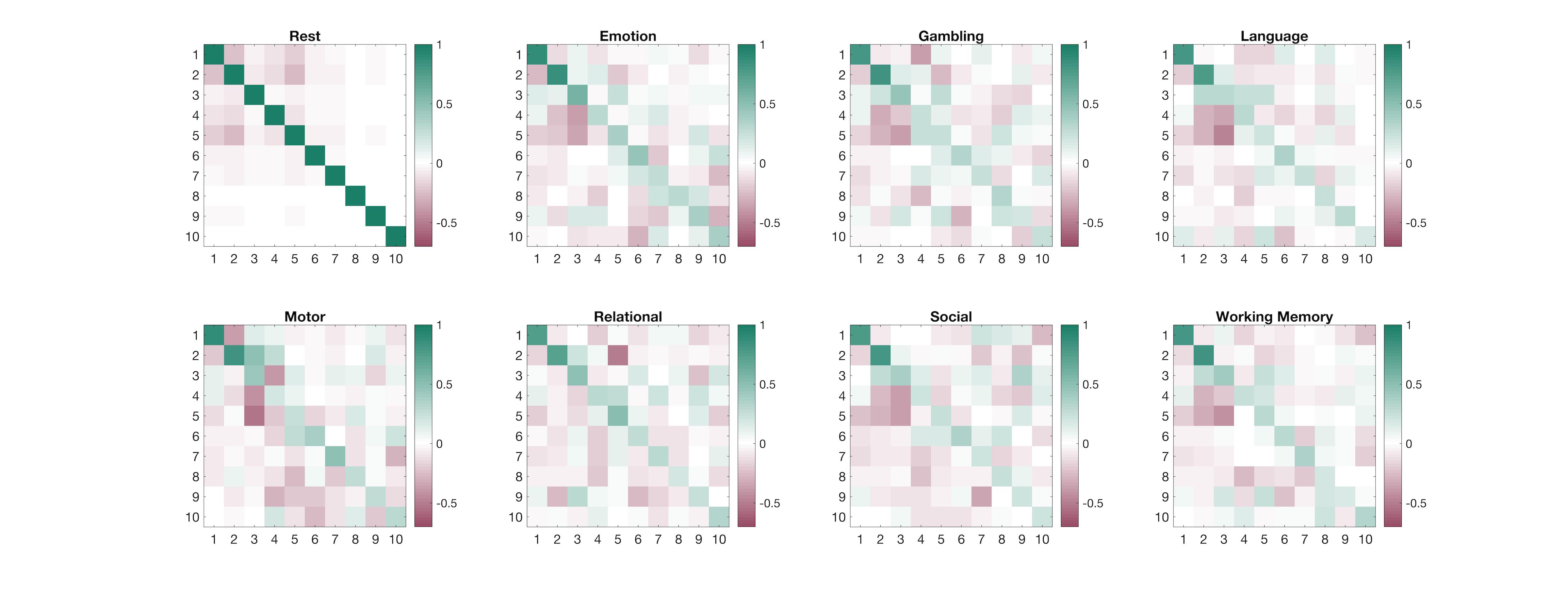

**Supplementary Figure S10**. Similarity of reordered task-related components with task-free components. Rows represent the first ten task-free components and columns represent the first ten task-related components or task-free (rest) components.
